# Next-generation mapping of the salicylic acid signaling hub and transcriptional cascade

**DOI:** 10.1101/2024.01.03.574047

**Authors:** Jordan Powers, Xing Zhang, Andres V. Reyes, Raul Zavaliev, Shou-Ling Xu, Xinnian Dong

**Affiliations:** Howard Hughes Medical Institute, Duke University, Durham, NC 27708, USA; University Program in Genetics and Genomics, Duke University, Durham, NC 27708, USA; Carnegie Institute for Science, Stanford University, Stanford, CA 94305, USA

## Abstract

For over 60 years, salicylic acid (SA) has been known as a plant immune signal required for both basal and systemic acquired resistance (SAR). SA activates these immune responses by reprogramming up to 20% of the transcriptome through the function of NPR1. However, components in the NPR1-signaling hub, which appears as nuclear condensates, and the NPR1-signaling cascade remained elusive due to difficulties in studying transcriptional cofactors whose chromatin associations are often indirect and transient. To overcome this challenge, we applied TurboID to divulge the NPR1-proxiome, which detected almost all known NPR1-interactors as well as new components of transcription-related complexes. Testing of new components showed that chromatin remodeling and histone demethylation contribute to SA-induced resistance. Globally, NPR1-proxiome shares a striking similarity to GBPL3-proxiome involved in SA synthesis, except associated transcription factors (TFs), suggesting that common regulatory modules are recruited to reprogram specific transcriptomes by transcriptional cofactors, like NPR1, through binding to unique TFs. Stepwise greenCUT&RUN analyses showed that, upon SA-induction, NPR1 initiates the transcriptional cascade primarily through association with TGA TFs to induce expression of secondary TFs, predominantly WRKYs. WRKY54 and WRKY70 then play a major role in inducing immune-output genes without interacting with NPR1 at the chromatin. Moreover, a loss of NPR1 condensate formation decreases its chromatin-association and transcriptional activity, indicating the importance of condensates in organizing the NPR1-signaling hub and initiating the transcriptional cascade. This study demonstrates how combinatorial applications of TurboID and stepwise greenCUT&RUN transcend traditional genetic methods to globally map signaling hubs and transcriptional cascades.

## INTRODUCTION

In plants, a local infection can often lead to systemic acquired resistance (SAR) through the accumulation of the phytohormone, salicylic acid (SA)^1^, which, in *Arabidopsis,* could result in changes in up to 20% of its transcriptome^2^. This process is mediated by the downstream signal component nonexpresser of PR genes 1 (NPR1); mutating *NPR1* leads to a drastic loss of this transcriptional response and enhanced susceptibility to primary and secondary infection^3^. Since the NPR1 protein lacks a DNA-binding domain, it is proposed to function as a transcriptional cofactor for transcription factors (TFs) such as TGAs^4–6^ and WRKYs^7,8^. However, our knowledge on how NPR1 functions molecularly to orchestrate the transcriptome-wide changes in response to SA is still limited by the insufficient sensitivity of current methodologies for investigating a transcriptional cofactor like NPR1. Recent structural study of the NPR1 complex with TGA3 TF showed that NPR1 serves its transcriptional coactivator role as a dimer by bridging two dimeric TGA3 molecules^9^. The presence of (NPR1)_2_-(TGA3)_2_ intermediates in the cryo-EM samples suggests that the NPR1 dimer may function as a platform to nucleate different TFs in an enhanceosome. This raises the question, does NPR1 interact with different TFs concurrently in response to SA to activate the myriad of output genes or, alternatively, initiate the reprogramming through a transcriptional cascade? Besides TFs, NPR1 is likely to be associated with large molecular complexes in response to SA because of the nuclear and cytoplasmic condensates detected for the protein^7,10^. While the cytoplasmic condensates (cSINCs) have been characterized^10^, the function and contents of NPR1 nuclear condensates (nSINCs) remain to be elucidated. Therefore, a comprehensive study of NPR1’s proximal partners in the nucleus and a stepwise dissection of NPR1 transcriptional targets are essential for elucidating the molecular mechanisms by which this master immune regulator reprograms the transcriptome.

## RESULTS

### Label-free quantitative analysis of NPR1-proxiome using TurboID led to identification of new regulators of SA-induced resistance

To address the question how the transcriptional reprogramming occurs after the SA-bound NPR1 dimer bridges the TGA TF complexes^9^, we generated stable transgenic plants expressing NPR1-3xHA fused with a promiscuous biotin ligase, TurboID^11,12^. The activity of the resulting NPR1-3xHA-TurboID (NPR1-TbID) was validated by its ability to restore, in the *npr1-2* background, the induction of *PR1*, a known NPR1 target (Extended Data Fig. 1a). Based on the *PR1* gene expression pattern, we treated the transgenic line with 1 mM SA for 4 h, when *PR1* is showing the most rapid increase^7^, followed by sample collection and processing under either a stringent or a harsh condition (see Methods). Using label-free quantification of the LC-MS/MS data^13^, we identified 234 NPR1-proximal proteins (FC_LFQ_ ≥ 2, p-value < 0.01 in either condition or p-value < 0.1 in both conditions) enriched in the NPR1-TurboID sample compared to the control (Fig. 1a, Extended Data Fig. 1b, Supplementary Data 1). For the first time, we were able to detect almost all known, key NPR1 interactors identified through decades of genetic and molecular studies, including NPR1-like protein 3 (NPR3) and 4 (NPR4)^14^, NIM1-interacting 1 (NIMIN1)^15^, TGA5^4–6^, WRKY18^8^, histone acetyltransferase of the CBP family 1 (HAC1)^16^, and components of Mediator^17^ (Fig. 1b), validating the superior sensitivity of the method. Critically, the identified proximal proteins have minimal overlap with the components of SA-induced NPR1 condensates in the cytoplasm, cSINCs^10^ (Fig. 1c), giving us confidence in the identification of the NPR1 nuclear proxiome, which are possible components of the NPR1-enhanceosome.

**Fig. 1.**
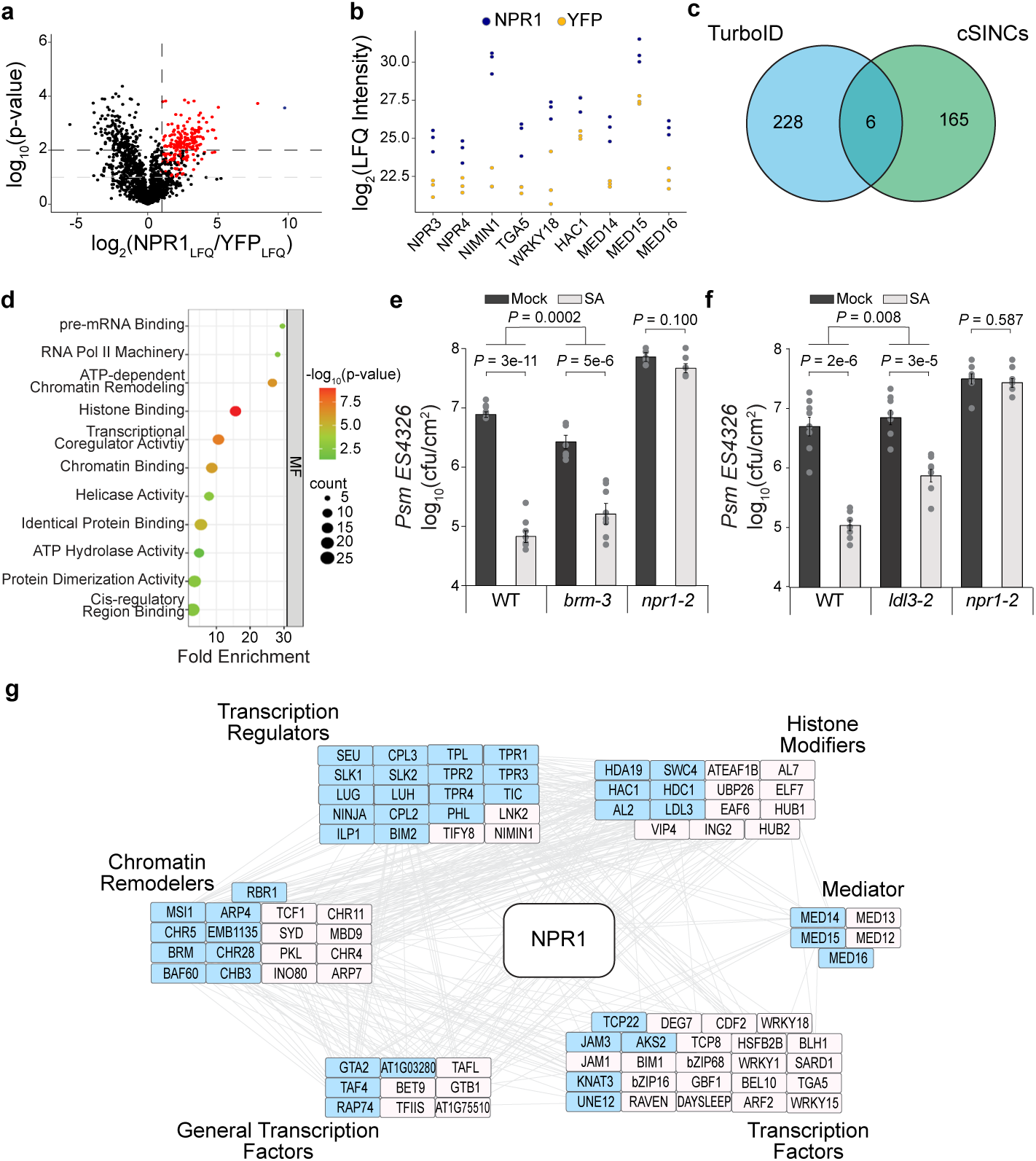
NPR1-proxiome contains transcriptional machineries and chromatin remodelers shared by GBPL3-proxiome. **a,** Volcano plot of NPR1 proximal proteins 4 h after SA treatment detected through TurboID biotin affinity purification followed by Label Free Quantification (LFQ) Mass Spectrometry processed under stringent conditions (see Methods). Red points represent proteins that have a NPR1_LQF_/YFP_LFQ_ ≥ 2 and p-value < 0.1 in both stringent and harsh washing conditions (see Methods) or p-value < 0.01 in at least one washing condition. The single blue point represents NPR1. **b,** Log_2_(Maximum LFQ Intensity) of known NPR1 interactors in NPR1-TbID (NPR1) vs. YFP-YFP-TbID (YFP) samples. **c,** Venn diagram comparing NPR1 proximal proteins identified in the current TurboID experiment with those identified in the cytoplasmic SA-induced NPR1 condensates (cSINCs)^10^. **d,** Enriched molecular functions (MF) of the 234 NPR1 proximal proteins. **e, f,** WT, *npr1-2*, *brm-3* (**e**), and *ldl3-2* (**f**) treated with H_2_O (mock) or 1 mM SA for 24 h prior to inoculation with *Psm* ES4326 at OD_600 nm_ = 0.001. Bacterial colony-forming units (cfu) were measured 3 days post inoculation (n = 8; error bars represent SEM; two-sided t-test and two-way ANOVA were used for comparisons within and between genotypes, respectively). **g,** STRING network analysis^43^ of NPR1 proximal proteins relating to chromatin remodeling and transcriptional regulation. Blue shade, proteins shared with GBPL3-proxiome^21^.

Excitingly, this analysis also identified many new NPR1 proximal partners. Gene Ontology (GO) term analysis based on molecular function (MF) demonstrated that these partners are enriched with proteins involved in histone modifications, chromatin remodeling, transcriptional machinery, and splicing complexes (Fig. 1d, Extended Data Fig. 1c), suggesting roles of these nuclear functions in reprogramming the SA transcriptome. The multi-functional feature of the NPR1-proxiome is consistent with its central role as a signaling hub for conferring disease resistance against a broad-spectrum of pathogens and abiotic stresses^10,18,19^. To begin to functionally validate the NPR1-proximal complexes, we focused on two groups of NPR1 partners: (1) the chromatin remodeling SWItch/Sucrose Non-Fermentable (SWI/SNF) proteins, with BRAHMA (BRM) as a representative, and (2) the histone modifying proteins, with the histone demethylase LSD1-like 3 (LDL3) as a representative. Although chromatin remodeling, nucleosome repositioning, and histone modifications have previously been shown to occur at SA-responsive genes and may play a role in their induction^16,20^, the involvements of BRM and LDL3 have not been tested in SA-induced resistance. We found that knocking out the *BRM* and *LDL3* genes partially compromised SA-induced resistance to the bacterial pathogen *Pseudomonas syringae* pv *maculicola* ES4326 (*Psm* ES4326) (Fig. 1e, f), indicating that chromatin remodeling through BRM and histone demethylation through LDL3 are important regulatory steps involved in SA/NPR1-mediated transcriptional reprogramming. The moderate phenotypes of these mutants highlight the effectiveness of TurboID in identifying new signaling mechanisms involved in essential and robust cellular processes which are difficult to uncover using forward genetic approaches due to the moderate phenotypes of the viable mutants.

Interestingly, both of BRM and LDL3 proteins have been reported in proximity with the condensate-forming protein, Guanylate-Binding Protein-Like 3 (GBPL3)^21^. Similar to GBPL3, which has been shown to be involved in temperature-mediated SA synthesis and pathogen response^22,23^, NPR1 also forms nuclear condensates in response to SA induction^7^. From a more in-depth comparison between the NPR1-proxiome and the GBPL3-proxiome, we discovered a large overlap in transcriptional regulators, chromatin remodelers, and histone modifiers (Fig. 1g, shaded in blue). However, most of the TFs appeared to be NPR1-specific partners (19/24). This led to an exciting hypothesis that plants reprogram their transcriptome in response to a specific stimulus by recruiting common transcriptional regulatory modules and machineries, but unique TFs, by a hub protein, such as NPR1, which has the intrinsic property to form biomolecular condensates^24^.

### QuantSeq shows WRKY54 and WRKY70 are positive regulators of SA/NPR1-mediated transcriptional reprogramming

Among the TFs unique to NPR1 based on our TurboID data, TGA and WRKY TFs have already been observed to interact with NPR1 in previous studies^4–8,10^(Fig. 1b, g). However, while TGA3 TF has been shown to bind DNA in complex with NPR1 in the cryo-EM structure^9^, the transcriptional role of WRKY TFs in SA-mediated gene expression is less straightforward because different WRKYs, with their own transcription induced by various stresses, have redundant and distinct roles in regulating gene expression^25^. In this study, we focused on WRKY70 and its closest homolog WRKY54 (WRKY54/70), because they have been shown to associate with NPR1^7^. We performed QuantSeq^26^ in WT, *npr1-2*, and *wrky54 wrky70* (*wrky54/70*) double mutants 8 h after SA induction and identified 2528 differentially expressed genes in response to SA (|log_2_foldchange| ≥ 1, adjusted p-value < 0.1), of which 1909 were induced and 1619 were repressed (Extended Data Fig. 2a, Supplementary Data 2). Among the 1909 SA-induced genes, 1022 were NPR1-dependent and 782 were WRKY54/70-dependent (Extended Data Fig. 2b, Supplementary Data 2), and the global transcriptome displayed a higher degree of correlation with NPR1 than with WRKY54/70 (Extended Data Fig. 2c, d). GO analyses of NPR1- and/or WRKY-dependent genes did not provide further resolution, with similar enrichments for defense response and SA-related processes (Extended Data Fig. 3a-c). Interestingly, promoter examination of these genes led to detection of the WRKY-binding “W-box” as the most enriched motif (Extended Data Fig. 3d-f), instead of the *as-1* element for TGA TFs, even for those NPR1-dependent, WRKY54/70-independent genes (Extended Data Fig. 3f), suggesting that WRKY TFs are the major TFs responsible for the SA-mediated transcriptional output.

**Fig. 2.**
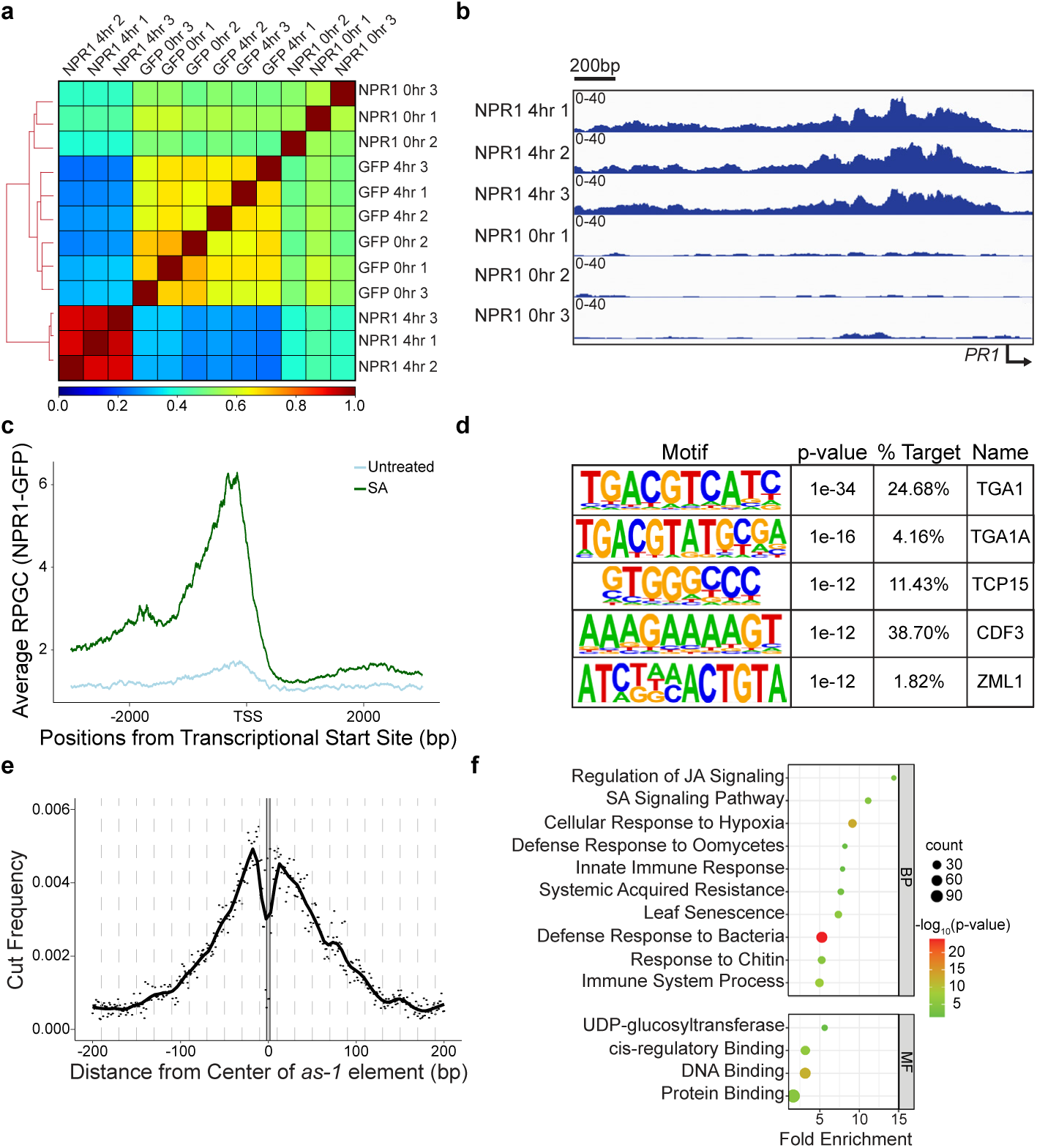
NPR1 directly targets TF genes through association with TGA TFs. **a**, Pearson’s correlation of the greenCUT&RUN data from plants expressing NPR1-GFP (NPR1) and GFP with and without 1 mM SA treatment for 4 h. **b,** Integrative Genomics Viewer (IGV) of the *PR1* promoter showing normalized NPR1-GFP binding before and after SA treatment. **c,** Mean profile of Reads Per Genomic Content (RPGC) of NPR1-GFP reads before and after SA treatment at NPR1-target genes. TSS, transcriptional start site. **d,** Motifs enriched under NPR1-GFP peaks. **e,** Cut frequency of all *as-1* element (TGACG) by the GFP nanobody-MNase in the overall NPR1-GFP peaks 4 h after 1 mM SA treatment. **f,** The enriched biological processes (BP) and molecular functions (MF) of NPR1-target genes.

**Fig. 3.**
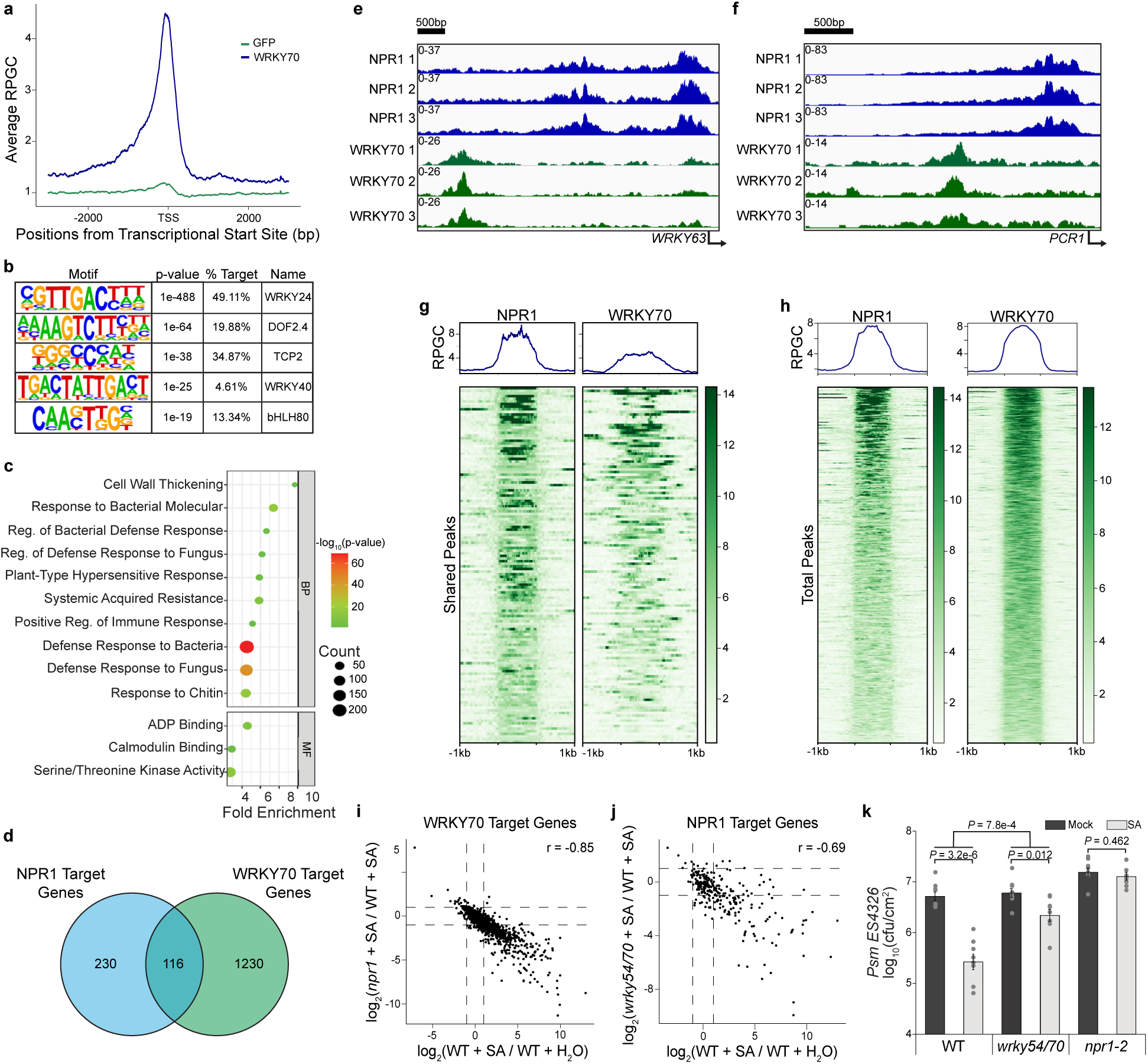
WRKY54/70 are major TFs downstream of NPR1-TGA that positively regulate SA-mediated gene expression. **a**, Mean profile of Reads Per Genomic Content (RPGC) of WRKY70-GFP (WRKY70) and GFP reads of WRKY70-target genes. TSS, transcriptional start site. **b,** Motifs enriched under WRKY70-GFP peaks. **c,** Enriched biological processes (BP) and molecular functions (MF) of WRKY70-target genes. **d,** Venn diagram illustrating the overlap between NPR1- and WRKY70-target genes. **e, f,** Integrative Genomics Viewer (IGV) of normalized NPR1 and WRKY70 binding at the promoters of their shared target genes *WRKY63* (**e**) and *PCR1* (**f**). **g,** RPGC of NPR1-GFP and WRKY70-GFP at 116 shared target genes 1 kb upstream and downstream of NPR1 peaks. **h,** RPGC of all NPR1-GFP and WRKY70-GFP target genes centered on their respective peaks. **i,** Correlation between SA-induced transcription and NPR1-dependency in WRKY70-target genes. r, Pearson correlation coefficient. **j,** Correlation between SA-induced transcription and WRKY54/70-dependency in NPR1-target genes. **k,** Bacterial colony-forming units (cfu) in WT, *wrky54/70*, and *npr1-2.* Plants were treated with H_2_O (mock) or 1 mM SA for 24 h before inoculated with *Psm* ES4326 at OD_600 nm_ = 0.001. CFUs were measured 2 days post inoculation (n = 8; error bars represent SEM; two-sided t-test and two-way ANOVA were used for comparison within and between genotypes, respectively).

### Genome-wide greenCUT&RUN identified WRKY TF genes as a major group of NPR1 transcriptional targets

The enrichment of the W-box in our QuantSeq data (Extended Data Fig. 3d-e) and in other transcriptome profiling datasets at various time points after SA or SA analog treatment^2,16,27,28^ (Extended Data Fig. 4a-c) raised the question about the role of TGA TFs in the SA signaling cascade and the relationship between TGA and WRKY TFs with NPR1. To address these questions, we performed Cleavage Under Target and Release Using Nuclease (CUT&RUN) followed by next generation sequencing^29^ 4 h after SA induction to identify direct transcriptional targets of NPR1, utilizing an anti-GFP antibody on *35S:NPR1-GFP* and *35S:GFP* transgenic plants. Unfortunately, the experiment failed to detect any differential peaks between NPR1-GFP and GFP samples with minimal difference seen at either known NPR1 targets or globally (Extended Data Fig. 5). This suggests that while CUT&RUN has significantly enhanced sensitivity for identifying TFs that interact directly with chromatin^30^ and histone modifications^31^, an even more sensitive methodology is required for detecting targets of transcriptional cofactors, like NPR1, whose proximity to DNA depends on its interaction with TFs.

**Fig. 4.**
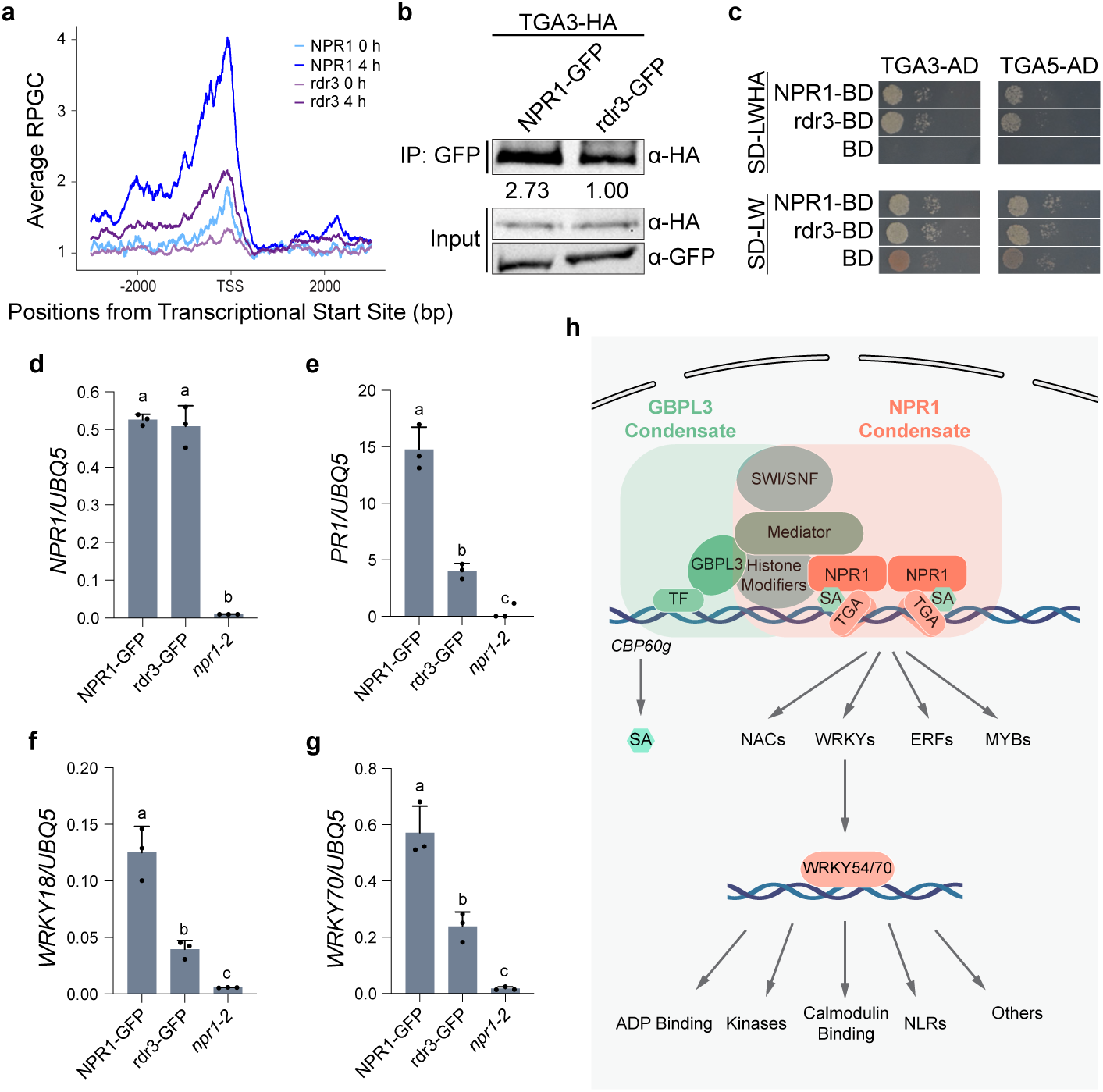
Biomolecular condensate formation stabilizes NPR1 association with TGA TF and enhances its transcriptional activity. **a**, Mean profile of Reads Per Genomic Content (RPGC) of NPR1-GFP (NPR1) and npr1^rdr3^-GFP (rdr3) 4 h after 1 mM SA treatment at NPR1-target genes. TSS, transcriptional start site. **b,** co-immunoprecipitation (co-IP) between TGA3 and NPR1 or rdr3 transiently overexpressed in *N. benthamiana*. Value under IP blot represents band intensities normalized to TGA3 input. **c,** Interaction between TGA3/TGA5 fused to the activator domain (AD) and NPR1/rdr3 fused to the DNA-binding domain (BD) in the yeast two-hybrid assay. Yeast strains were mated for 24 h, normalized to OD_600 nm_ = 1.0, serial diluted, plated on the indicated Synthetic Defined (SD) media without leucine and tryptophan (LW) or without leucine, tryptophan, histidine, and adenine (LWHA), and incubated at 30 °C. Photos were taken 2 days after plating. **d-g,** Transcript levels of *NPR1/rdr3* (**d**) and target genes *PR1* (**e**), *WRKY18* (**f**), and *WRKY70* (**g**) in *35S:NPR1-GFP/npr1-2*, *35S:npr1^rdr3^-GFP/npr1-2,* and *npr1-2* plants measured using qPCR 8 h after SA induction (n = 3, error bars represent standard deviation). **h,** Working model of the SA/NPR1 signaling hub and transcriptional cascade. Overlapped rectangles show that NPR1- and GBPL3-condensates share general transcriptional regulatory machineries (e.g., Mediator, SWI/SNF, and histone modifiers), but target different genes through association with unique TFs. An increase in SA level triggers the transcriptional cascade by first activating NPR1 to induce TGA-mediated expression of WRKY, MYB, NAC and ERF TFs which in turn activate the subsequent gene expression.

To further improve the sensitivity of the CUT&RUN methodology, which relies on transient interactions of multiple proteins that ultimately lead to the cutting of target DNA sequences by pA-MNase, we adopted an anti-GFP nanobody-based CUT&RUN approach, ‘greenCUT&RUN’, where the GFP-specific nanobody is fused directly to the MNase^32^. In contrast to the initial CUT&RUN data (Extended Data Fig. 5), the new method led to a clear separation of the SA-treated NPR1-GFP samples from both the untreated NPR1-GFP and the GFP samples (Fig. 2a). Based on the three NPR1-GFP replicates, we were able to detect 385 reproducible NPR1-GFP-specific peaks (Extended Data Fig. 6a, Supplementary Data 3). Furthermore, examining the promoter of the known NPR1 target gene, *PR1*, an SA-dependent accumulation of NPR1-GFP could be observed clearly (Fig. 2b). By averaging global alignment of the binding loci, we detected a significant enrichment of NPR1-GFP at the promoters of its target genes upon SA treatment compared to the untreated samples (Fig. 2c). Among these loci, 84.2% occurred upstream of the transcriptional start site (TSS). Interestingly, the distances from TSS of these binding peaks varied widely from gene to gene, ranging from immediately before the TSS to several thousand base pairs (kb) upstream, with only 53% within 1 kb from TSS (Extended Data Fig. 6b). These results are consistent with NPR1-enhanceosome’s function in bringing in distal binding sites through DNA loop and in recruiting larger transcriptional machineries like the SWI/SNF complex and Mediator^33,34^(Fig. 1g).

Among the NPR1 peaks, we detected the TGA-binding *as-1* element, TGACG, as the most significantly enriched motif (Fig. 2d). While there was an increased cutting frequency near the motif, the motif itself was protected from the MNase, further supporting the notion that NPR1 binds to the DNA through TGA TFs (Fig. 2e). Additionally, we also detected enrichment of Teosinte branched 1/Cycloidea/Proliferating cell factors (TCP) and Cycling Dof Factor (CDF) binding motifs (Fig. 2d), which are two other TFs detected in our TurboID experiment (Fig. 1g). As expected, the NPR1-target genes are largely related to defense response and hormone cross talk between SA and another plant defense hormone, jasmonic acid (JA) (Fig. 2f). Surprisingly, the W-box, the most enriched motif among SA-induced genes, was not enriched under NPR1 peaks. Moreover, compared to the thousands of differentially expressed genes in response to SA, there were only a few hundred NPR1-target genes. These data suggest that NPR1 reprograms the transcriptome through multiple steps, instead of through parallel association with multiple TFs. In support of this hypothesis, the GO terms of NPR1 transcriptional targets are largely enriched with TFs and other DNA binding proteins (Fig. 2f). Analysis of the genes annotated as DNA binding and/or cis-regulatory binding detected four major TF families: WRKYs, NACs, ERFs, and MYBs, with WRKYs representing the largest family (Extended Data Fig. 7a). Of note, NPR1 preferentially targets group III WRKY TFs, including WRKY70 (Extended Data Fig. 7b, c), suggesting their involvement in further propagating SA-induced gene expression.

### Genome-wide greenCUT&RUN established WRKY70 as a downstream TF in the SA-induced transcriptional cascade

To examine the role of group III WRKYs in SA/NPR1-mediated reprogramming of the immune transcriptome, we performed a subsequent greenCUT&RUN analysis on a transgenic line *35S:WRKY70-GFP*. We collected the samples 2 h after SA treatment to take into consideration of the previous hypothesis that WRKY70 repression on the marker gene *PR1* is removed by NPR1 prior to its activation of TGA TF^7^. Similar to our NPR1 greenCUT&RUN experiment, we found that WRKY70-GFP samples were well-correlated with one another, while distinguished from those of the GFP samples. Surprisingly, they were also distinct from the NPR1-GFP greenCUT&RUN data (Extended Data Fig. 8a). From this experiment, we detected 1477 reproducible WRKY70-GFP-specific peaks (Extended Data Fig. 8b, Supplementary Data 4). It was evident that the WRKY70-GFP samples had a higher percentage of reproducible peaks (43.4% - 61.3%) compared to those in the NPR1-GFP samples (32.3% - 36.3%) (Extended Data Figs. 6a and 8b), consistent with WRKY70 being a TF. Examining all target genes showed that WRKY70, like NPR1, was mainly detected at the promoters of its target genes with only 14.4% of WRKY70 >1 kb upstream of TSS compared to the 31.2% for NPR1 (Extended Data Figs. 6b and 8c). As expected, a high enrichment of W-box was observed in these WRKY70-bound loci (Fig. 3b). Interestingly, while defense-related biological processes were still the top enrichments in the WRKY-target genes, they differ from those of NPR1-target genes in their molecular functions. Where NPR1 targets TF genes, WRKY70 targets those involved in ADP-binding (mostly encoding nucleotide-binding domain and leucine-rich repeat-containing immune receptors, NB-LRRs), calmodulin-binding, and kinase activity (Fig. 3c), implying that WRKY70, whose transcription is induced by NPR1-TGA^2^ (Extended Data Fig. 7b), is involved in the downstream events in the signaling cascade of NPR1-mediated transcriptional reprogramming.

Apart from these distinct transcriptional targets, there were a smaller number of shared target genes between WRKY70 (116/1476) and NPR1 (116/346) (Fig. 3d), suggesting a possible interplay between WRKY70 and NPR1 in regulating the transcription of these genes. Investigating individual peaks, we saw WRKY70 and NPR1 indeed target the promoter of the same genes, but at distinct loci from one another (Fig. 3e, f). Interestingly, *PR1* was not detected in our WRKY70- GFP samples, despite the negative regulation WRKY70 has on the transcript^35^. Interestingly, when examining the peak patterns at all the shared target gene promoters, NPR1 samples showed one distinct peak (Fig. 3g), typical of its targets (Fig. 3h), while WRKY70 samples displayed much more varied and spreading peaks (Fig. 3g), which is atypical for the majority of the WRKY70 targets, where little spread is detected outside of the peak region (Fig. 3h). These data demonstrate that NPR1 is unlikely to switch association between WRKY and TGA TFs at the chromatin level as previously proposed^7^. Instead, NPR1 has been found to interact with WRKY70 in the cytoplasmic SINCs to sequester and degrade it^10^. Nevertheless, the shared gene targets of NPR1 and WRKY70 with distinct loci suggest a possible regulatory dependence on both proteins.

The sequential NPR1- and WRKY70-greenCUT&RUN analyses elucidated an SA-signaling cascade in which the SA-activated NPR1 induces the expression of *WRKY* TF genes through association with TGA TFs. Consistently, by comparing our QuantSeq results with NPR1- and WRKY70-greenCUT&RUN targets, we found that, while NPR1 had the expected strong regulation of WRKY70-target genes (r = 0.85) (Fig. 3i), WRKY54 and WRKY70 also had a moderate correlation with NPR1-targets (r = 0.69) (Fig. 3j), suggesting that, in addition to their role as feedback repressors of SA synthesis^2^, WRKY54 and WRKY70 are predominantly positive TFs of SA-mediated gene transcription. This hypothesis is further supported by the compromised SA-mediated resistance to *Psm* ES4326 observed in the *wrky54/70* double mutant compared to WT (Fig. 3k).

### SA-induced condensate-formation of NPR1 promotes its binding to the chromatin and transcriptional activity

With the identification of NPR1 proximal partners and direct transcriptional targets in the signaling cascade, we then tested our hypothesis that SA-induced condensate formation is critical for NPR1 to organize the enhanceosome to initiate transcription. We first performed greenCUT&RUN in the npr1^rdr^^3^-GFP mutant (referred to as rdr3)^10^, which can still translocate into the nucleus upon SA induction, but fails to form either nuclear or cytoplasmic condensates^10^. We found that chromatin association of rdr3 was still dependent on SA and occurred at the same loci as the WT NPR1, but at a significantly lower level (Fig. 4a), despite the fact that the mutant protein has a higher-than-WT nuclear distribution^10^. Interestingly, the reduced rdr3 binding to the TGA TF was only observed *in planta* (Fig. 4b), not in the yeast two-hybrid assay (Fig. 4c), suggesting that the decreased rdr3-chromatin association is less likely due to its diminished binding to TGA3 than the reduced stability of its complex with TGA3 due to inability to form the nuclear condensates. Moreover, at the same transcript levels (Fig. 4d), rdr3 had significantly compromised activity in inducing the direct target genes, *PR1* (Fig. 4e), *WRKY18* (Fig. 4f), and *WRKY70* (Fig. 4g) compared to the WT NPR1 control, supporting our hypothesis that NPR1 orchestrates the transcriptomic changes upon SA-induction by forming biomolecular condensates.

## DISCUSSION

By combinatorial applications of label-free quantification of TurboID-based LC-MS/MS data and the greenCUT&RUN technology, we have transcended, in a single study, decades of molecular genetic studies to generate a comprehensive map of the NPR1-centered transcriptional reprogramming machineries and the transcriptional cascade in response to SA induction (Fig. 4h). The validation of the new NPR1 proximal partners (Fig. 1e, f) clearly demonstrates the effectiveness of the methodology in studying signaling hubs formed by proteins, like NPR1, in association with regulatory modules involved in common nuclear functions, such as chromatin remodeling, histone modifications, Mediator, and RNA splicing. The robustness of these essential cellular machineries makes it difficult to discern their contributions to specific biological processes through genetic studies. Indeed, the NPR1-proxiome shows high similarity to the GBPL3- proxiome^21^, with the major distinction mainly in their associated TFs (Fig. 1g). Since both the GBPL3-proxiome required for inducing SA synthesis upon stress^22^ and the NPR1-proxiome responsible for SA-mediated transcriptional reprogramming can form nuclear biomolecular condensates^7^, it is tempting to hypothesize that in the nucleus, a similar set of transcriptional regulatory modules are recruited to form supramolecular complexes/condensates by distinct regulators, like NPR1, whose association with unique TFs provides the complexes/condensates functional specificity (Figs. 1g and 4h). Furthermore, condensate formation facilitates NPR1’s association with the chromatin, as well as target gene induction (Fig. 4a, d-g), supporting the notion that SA-induced nuclear NPR1-condensates, i.e., nSINCs, are transcriptionally active.

More experiments are required to demonstrate that NPR1 condensate formation is required for the recruitment of the transcriptional regulatory modules identified in the NPR1-proxiome (Fig. 1a, Extended Data Fig. 1b, Supplementary Data 1). In the survey of genome-wide association of the key chromatin remodeling protein BRM using a stable *BRM:BRM-GFP* transgenic line^36^, we found that while the overall BRM-specific peaks stayed constant under both mock and SA-induced conditions, indicating that SA has minimal impact on the general BRM binding to the chromatin (Extended Data Fig. 9a, Supplementary Data 5), its association to the NPR1-targeted loci was enhanced by SA treatment (Extended Data Fig. 9b), suggesting that BRM is recruited to NPR1-target genes upon induction. However, the significant basal levels of BRM at these NPR1 loci before SA induction indicate that members of this transcriptional machinery may already be present at the target gene promoters. It would be exciting to explore which proteins of these transcriptional modules are constitutively present at the promoters and which are recruited in response to induction to initiate transcription. Consistent with NPR1 condensate formation being a dynamic process, SA/NPR1-induced WRKYs as well as several known negative regulators of SA-mediated gene expression, such as NPR3, NPR4, NIMINs, and TPLs, were found to be in the NPR1-proxiome. However, we cannot rule out the possibility that the NPR1-proxiome consists of multiple distinct NPR1-protein complexes. Future research will be required to understand the dynamics of the NPR1 signaling hub.

Our success in using stepwise greenCUT&RUN to detect NPR1 direct targets and elucidating the hierarchical relationship between TGA and WRKY TFs demonstrates the method’s great potential in dissecting transcriptional cascades by providing a higher resolution than other transcriptomic methods tested in the study. As shown in our QuantSeq experiments with WT, *npr1* and *wrky54/70* mutants, the initiation step of the SA signaling cascade by NPR1 through TGA TFs was obscured because NPR1/TGA-targets were out-numbered by the subsequent WRKY-mediated transcriptional targets in the statistical analyses. Moreover, transcriptomic studies of TF gene families often rely on the usage of available TF knockdown lines or knockout mutants, which either have weak phenotypes due to functional redundancy or pleiotropic defects when higher order mutants are used. These limitations can now be overcome by the greenCUT&RUN method, which is readily applicable for studying not only TFs, but also any protein with indirect chromatin association.

## METHODS

### Plant material and growth conditions

All plants used in this study were grown on soil (ProMix B) under 12-h light/12-h dark conditions. The *35S:YFP-YFP-TbID* line was generously gifted by Dr. Zhi-Yong Wang^37^. The *35S:NPR1-3xHA-TbID* and *35S:npr1^rdr^*^3^*-GFP* constructs were made using Gateway cloning (Thermo Fisher Scientific). *35S:NPR1-3xHA-TbID* was transformed into the *npr1-2* plants using the floral dip method^38^. The *brm-3* (SALK_088462) and *ldl3-2* (SALK_146733) mutants were obtained from ABRC. The *35S:NPR1-GFP*, *35S:npr1^rdr^*^3^*-GFP*, and *35S:WRKY70-GFP* transgenic lines and the *wrky54 wrky70* double mutant were previously described^2,10^. The *BRM:BRM-GFP* line was a generous gift from Dr. Chenlong Li^36^.

### RNA isolation and qPCR

Total RNA was extracted from 3-week-old plants treated with 1 mM SA or H_2_O using Trizol^39^ (Thermo Fisher Scientific). DNase-treated total RNA was then used for SuperScriptIII Reverse Transcription (Thermo Fisher Scientific). The resulting cDNA samples were diluted tenfold for qPCR reactions using SYBR Green Master Mix to detect transcript levels.

### Affinity purification of biotinylated proteins

Affinity purification of biotinylated proteins was performed as previously described^11^, with minor modifications. Briefly, three replicates (4 g/sample) of 3-week-old plants treated first with 1 mM SA and, 1 h later, with 50 μM biotin for 3 h, were collected, flash frozen, and stored at -80 °C. Samples were ground to a fine powder, dissolved in 4 mL of the extraction buffer (50 mM Tris-HCl pH 7.5, 150 mM NaCl, 0.1% SDS, 1% NP-40, 0.5% Na-deoxycholate, 1 mM EGTA, 1 mM DTT, and the protease inhibitor cocktail), filtered, and sonicated. Sonicated samples were centrifuged, and biotin was removed from the resulting protein solution using PD-10 desalting columns (GE-Healthcare). The flow-through was collected and subjected to affinity purification using the streptavidin bead (Thermo Fisher Scientific). The resulting samples on the streptavidin beads were processed under two conditions: harsh and stringent. The harsh condition involved washing the beads 2x with the extraction buffer, 1x with 1 M KCl, 1x with 100 mM Na_2_CO_3_, 1x with 2 M Urea in 10 mM Tris-HCl pH 8, and 2x again with the extraction buffer. The stringent condition involved washing the beads 7x with the extraction buffer. The processed beads from both conditions were resuspended in 1 mL of the extraction buffer for further processing. Prior to trypsin digestion, the beads underwent further washes. The bead samples corresponding to the harsh conditions were followed by harsh washes consisting of 1x with cold 1 M KCl, 1x with 2 M Urea in 10 mM Tris-HCl pH 8, 2x with cold 50 mM Tris-HCl pH 7.5, and 2x with the Urea wash buffer (50 mM Tris-HCl pH 7.5, 1 M Urea). The bead samples corresponding to the stringent conditions were followed by mild washes consisting of 7x with the PBS buffer. Both sample sets were subjected to a 3 h incubation in 80 µl Trypsin buffer (50 mM Tris-HCl pH 7.5, 1 M Urea, 1 mM DTT, and 0.4 µg Trypsin) at 25 °C. The supernatants from the tryptic digest were transferred to new tubes and the beads were washed 2x with 60 µl 1 M Urea in 50 mM Tris-HCl pH 7.5. The combined 200 µL elutes were reduced (final concentration of 4 mM DTT), alkylated (final concentration of 10 mM Iodoacetamide), and digested overnight with 0.5 µg Trypsin. Additional 0.5 μg of trypsin was added in the next morning followed by acidification 4 h later by adding formic acid to a final concentration of ∼ 1 % and desalting using OMIX C18 pipette tips (A57003100).

### LC-MS/MS

LC-MS/MS was carried out on a Q-Exactive HF hybrid quadrupole-Orbitrap mass spectrometer (Thermo Fisher Scientific), equipped with an Easy LC 1200 UPLC liquid chromatography system (Thermo Fisher Scientific). Peptides were first trapped using a trapping column (Acclaim PepMap 100 C18 HPLC, 75 μm particle size, 2 cm bed length), then separated using analytical column AUR2-25075C18A, 25CM Aurora Series Emitter Column (25 cm x 75 µm, 1.6 μm C18) (IonOpticks). The flow rate was 300 nL/min, and a 120-min gradient was used. Peptides were eluted by a gradient from 3 to 28% solvent B (80% acetonitrile, 0.1% formic acid) over 100 min and from 28 to 44% solvent B over 20 min, followed by a 10 min wash at 90% solvent B. Precursor scan was from mass-to-charge ratio (m/z) 375 to 1,600 and top 20 most intense multiply charged precursors were selected for fragmentation. Peptides were fragmented with higher-energy collision dissociation (HCD) with normalized collision energy (NCE) 27.

### Proteomic analysis

Harsh and stringent sets of LC-MS/MS spectra were searched separately against the Araport11 database (20220914 version containing 49,467 entries) using the MSFragger 3.2^40^ software under default criteria to obtain maximum Label Free Quantification (LFQ) intensities. The search results were analyzed separately in Perseus^41^ (version 1.6.15.0). The processing in Perseus was as follows: MaxLFQ intensities were log2 transformed. Only proteins that had at least two valid values in at least one group (NPR1-TbID or YFP-YFP-TbID) were kept. The remaining missing MaxLFQ intensities were then imputed from a normal distribution that is downshifted by 1.8 and a width of 0.3 column wise. A two-sample t-test was conducted with a permutation-based (n = 250) FDR = 0.01 and the S0 = 2. Significant NPR1 proximal partners were identified by the following criteria: (1) a p-value < 0.1 in both processing conditions and a NPR1_LFQ_/YFP_LFQ_ ≥ 2 or (2) a p-value < 0.01 in either processing condition and a NPR1_LFQ_/YFP_LFQ_ ≥ 2. GO Term Analysis was performed using PANTHER^42^. Interaction network was performed using STRING^43^. Plots were generated with ggplot2^44^, Cytoscape^45^, and SRplot.

### SA-induced resistance against bacterial infection

SA-induced resistance was measured as previously described^46^. Briefly, *Pseudomonas syringae* pv. *maculicola* ES4326 (*Psm* ES4326) was grown at 30 °C on plates containing the King’s B medium (KB) for 48 h before resuspended in 10 mM MgCl_2_. 3-week-old plants were pretreated with 1 mM SA or H_2_O for 24 h prior to infection with *Psm* ES4326 at OD_600 nm_ = 0.001. Leaf discs from 8 infected plants were collected 2 days (for *wrky54 wrky70*) or 3 days (for *brm-3* and *ldl3-2*) post infection and individually ground in 0.5 mL of 10 mM MgCl_2_, serially diluted, and plated on the KB medium supplemented with 100 μg/mL of streptomycin. Colonies were counted two days later.

### QuantSeq and data analysis

Total RNA was extracted from 3-week-old leaves treated with 1 mM SA or H_2_O for 8 h using Split RNA Extraction Kit (Lexogen GmbH). RNA concentration was measured with Qubit RNA BR assay (Thermo Fisher Scientific) and integrity was checked with Agilent 2100 Bioanalyzer. Approximately 400 ng of RNA was used for library construction using the QuantSeq 3’ mRNA Seq Library Prep FWD Kit for Illumina (Lexogen GmbH)^26^. All libraries were sequenced at 100 bp single-end reads using the Illumina system NextSeq1000. Raw reads were trimmed to 50 bp using Trim Galore^47^ and mapped to the TAIR10 genome using the STAR aligner^48^ under the Lexogen recommended parameters. Differential expression between SA- and H_2_O-treated samples was detected using DESeq2^49^ with an adjusted p-value < 0.1 and a fold-change ≥ 2. GO Term Analysis was performed using PANTHER^42^ and *de novo* motif enrichment was uncovered using HOMER^50^ by analyzing promoters of differentially expressed genes from 1000 bp upstream to 200 bp downstream of the transcriptional start sites.

### greenCUT&RUN

Six leaves from two plants were collected before and after treatment with 1 mM SA for 4 h and stored at -80 °C. Frozen samples were ground to a fine powder and dissolved in 15 mL of the lysis buffer (20 mM Tris-HCl pH 7.5, 20% glycerol, 20 mM KCl, 2 mM EDTA, 2.5 mM MgCl_2_, 8.56% sucrose, and the protease inhibitor cocktail). Samples were filtered sequentially through a 70-μm filter and a 40-μm filter before centrifuged at 1,500 x g at 4 °C for 10 min. The pellet was resuspended in the nuclei isolation buffer (20 mM Tris-HCl pH 7.5, 20% glycerol, 2.5 mM MgCl_2_, 0.2% Triton X-100, and the protease inhibitor cocktail) and centrifuged at 1,500 x g at 4 °C for 10 min. The above resuspension and centrifugation steps were repeated 4x, until the pellet was free of any green color. The pellet was resuspended in 1 mL of the greenCUT&RUN wash buffer (20 mM HEPES-KOH pH 7.5, 150 mM NaCl, 0.5 mM Spermidine, and the protease inhibitor cocktail). Isolated nuclei were then bound to 40 μL of Concanavalin A beads resuspended in 10 μL of binding buffer (20 mM HEPES-KOH pH 7.5, 10 mM KCl, 1 mM CaCl_2_, 1 mM MnCl_2_, and the protease inhibitor cocktail) and rotated for 10 min at room temperature. The beads were collected using a magnetic rack, the supernatant was then removed, the bound nuclei were then resuspended in 1 mL of the EDTA buffer (20 mM HEPES-KOH pH 7.5, 150 mM NaCl, 0.5 mM Spermidine, 2 mM EDTA, and the protease inhibitor cocktail), and rotated at room temperature for 10 min. The beads were collected again and resuspended in 100 μL of the greenCUT&RUN wash buffer containing 10 μg/mL of nanobody-MNase and rotated at 4 °C for 30 min. After rotation, beads were collected and washed twice in the greenCUT&RUN wash buffer. Beads were then put on ice, resuspended in 150 μL of the calcium buffer (20 mM HEPES-KOH pH 7.5, 150 mM NaCl, 0.5 mM Spermidine, 3 mM CaCl_2_, and the protease inhibitor cocktail) and incubated on ice for 30 min. After incubation, 100 μL of the 2X stop buffer (340 mM NaCl, 20 mM EDTA, 10 mM EGTA, 100 μg/mL RNase A, and 50 μg/mL Glycogen) was added to the beads and incubated at 37 °C for 30 min. After incubation, beads were removed, and the supernatant was collected for DNA isolation. 2 μL of 10% SDS and 20 μg of Proteinase K were added to the collected supernatant and incubated at 50 °C for 1 h. Equal volume of Phenol:Chloroform:Isoamyl Alcohol (25:24:1, v/v) was added to the samples followed by vortexing. The solution was transferred to a phase lock tube and centrifuged for 5 min at 16,000 x g at room temperature. After centrifugation, equal volume of chloroform was added, samples were inverted 10x, and centrifuged for 5 min at 16,000 x g at room temperature. The top aqueous layer was then taken and moved into new tubes containing 3 μL of 2 mg/mL of glycogen. 2x volumes of 100% ethanol was added to each sample to facilitate DNA precipitation overnight at -20 °C. After DNA precipitation, samples were centrifuged for 10 min at 16,000 x g at 4 °C. The supernatant was removed, the pellet was washed in 1 mL of 100% ethanol, and centrifuged for 5 min at 16,000 x g at 4 °C. The supernatant was removed, and the pellet was air dried for 5 to 10 min. The pellet was resuspended in 50 μL of H_2_O and used for library preparation.

### CUT&RUN

The protocol for nuclei isolation for the CUT&RUN protocol was the same as for greenCUT&RUN described above. After nuclei isolation, the previously reported CUT&RUN protocol^29^ was followed.

### Sequencing library construction for CUT&RUN and greenCUT&RUN

CUT&RUN and greenCUT&RUN libraries were constructed using the KAPA HyperPrep Kit (Roche Holding AG), with minor modifications. Briefly, end repair and A-tailing were performed at 20 °C for 30 min followed by deactivation of the A-tailing enzyme at 58 °C for 1 h. 1/100 diluted Illumina TruSeq DNA UD Indexes were ligated on to A-tailed DNA at 20 °C for 30 min. Post-ligation cleanup was performed twice, first using 1x library volume of AMPure Beads, next with 1.2x library volume of AMPure Beads, followed by a double-sided size selection to remove larger DNA fragments and smaller adapter dimers, respectively, using 0.7X-1.2X library volume of AMPure Beads following the manufacture’s protocol (Roche Holding AG). Ligated libraries were then amplified using PCR and cleaned up twice with 1.2x library volume of AMPure Beads to generate final purified libraries. Library size and concentration were determined using Agilent 2100 Bioanalyzer and Qubit (Thermo Fisher Scientific), respectively. The *35S:NPR1-GFP*, *35S:npr1^rdr^*^3^*-GFP*, *35S:WRKY70-GFP*, and *35S:GFP* (control) libraries were sequenced at 75 bp paired-end reads using the Illumina system NextSeq500. The *BRM:BRM-GFP* and *35S:GFP* (control) libraries were sequenced at 100 bp paired-end reads using the Illumina system NextSeq1000.

### CUT&RUN and greenCUT&RUN data analysis

Raw reads were trimmed using Trim Galore^47^ and aligned to the TAIR10 genome using bowtie2^51^. Concordant read Sequence Alignment Map (SAM) files were converted to Binary Alignment Map (BAM) files and PCR-duplicated reads were removed using SAMtools^52^. Deduplicated BAM files were then used to call peaks using MACS2^53^. Peaks called in all samples were used for further analysis. Bigwig and bedgraph files of normalized Read Per Genomic Content (RPGC) were generated using bamCoverage from deepTools 3.5.1^54^. Bigwig files were visualized in IGV^55^. Normalized bigwig files and deepTools 3.5.1 were used for generating Pearson correlation heatmaps and peak heatmaps. *De novo* motif prediction of reproducible peaks was performed using HOMER^50^. GO Term Analysis was performed using PANTHER^42^. Cut frequency plot was generated using cut-frequency^56^. Mean profile plots were generated using custom code in R.

### Yeast two-hybrid

AH109 and Y187 yeast strains were transformed with the TGAs/pGADT7 and NPR1s/pGBKT7 constructs, respectively. NPR1 and npr1^rdr^^3^ were used as the bait and TGA3 and TGA5 were used as the prey. All protocols were carried out according to Clontech Yeast Protocols Handbook.

### Protein analysis and immunoprecipitation (IP)

Protein analysis and IP were performed as previously described^57^. Briefly, recombinant proteins were transiently overexpressed in *Nicotiana benthamiana* by coinjecting the *Agrobacterium tumefaciens* strain GV3101 carrying the *35S:NPR1-GFP* construct (OD_600 nm_ = 0.5) or *35S:npr1^rdr^*^3^*-GFP* construct (OD_600 nm_ = 0.5) with the *Agrobacterium tumefaciens* strain GV3101 carrying the *35S:TGA3-HA* construct (OD_600 nm_ = 0.5) into the abaxial side of the leaf. After 44 h, plants were sprayed with 1 mM SA for 4 h before 1 g of tissue was collected and flash frozen. Frozen tissue was then ground and resuspended in 2.5 mL of the IP Buffer (10% glycerol, 25 mM Tris-HCl pH 7.5, 1 mM EDTA, 150 mM NaCl, 10 mM DTT, the protease inhibitor cocktail, and 0.2% NP-40). 40 mL of α-GFP beads (Chromotek) were added to the lysate for protein binding overnight at 4 °C, followed by 3x washes in the IP buffer. 50 μL of 4x LDS Sample Buffer (Thermo Fisher Scientific) was added to the beads and incubated at 70 °C for 20 min. Samples were then run on a 4-12% Bis-Tris gel and transferred to a membrane for western blotting using α-GFP (Clontech) and α-HA (Cell Signaling Technology) antibodies. Band intensity was measured using the iBright Analysis Software (Thermo Fisher Scientific).

### Statistics and reproducibility

In all statistical data, the center values are the mean, and the error bars all represent standard error of the mean except in Figure 4 qPCR data (standard deviation). All experiments were performed three or more times with similar results except Affinity Purification LC-MS/MS (once), QuantSeq (once), and greenCUT&RUN (once).

## ACKNOWLEDGEMENTS

We thank Dr. Marc Timmers for providing the GFP nanobody-fused MNase enzyme and the recombinant plasmid encoding the fusion protein; Dr. Zhi-Yong Wang for the gift of the YFP-YFP-TurboID line; Dr. Chenlong Li for sharing the *BRM:BRM-GFP* line; and members of the Dong laboratory for helpful discussion of the project. We thank the Duke University School of Medicine for the use of the Sequencing and Genomic Technologies Shared Resource for providing NGS services. This work was supported by grants from the National Institutes of Health (NIH) 1R35GM118036 and the Howard Hughes Medical Institute to X.D; NIH 5T32GM007754-40 to J.P., NIH R01GM135706 to S.-L.X. and its diversity supplement to support A.V.R, as well as the Carnegie endowment to the Carnegie mass spectrometry facility.

## AUTHOR CONTRIBUTIONS

X.D. conceived and supervised this project. J.P. carried out all the TurboID, CUT&RUN, greenCUT&RUN, QuantSeq, and molecular genetic experiments and performed the associated computational analyses. X.Z. generated the NPR1-3xHA-TurboID construct. A.R. and S.X. performed the LC-MS/MS of the TurboID samples. R.Z. carried out the marker gene expression analysis. J.P. and X.D. wrote the manuscript with input from all coauthors.

## Extended Data Figures and Figure Legends

**Extended Data Fig. 1.**
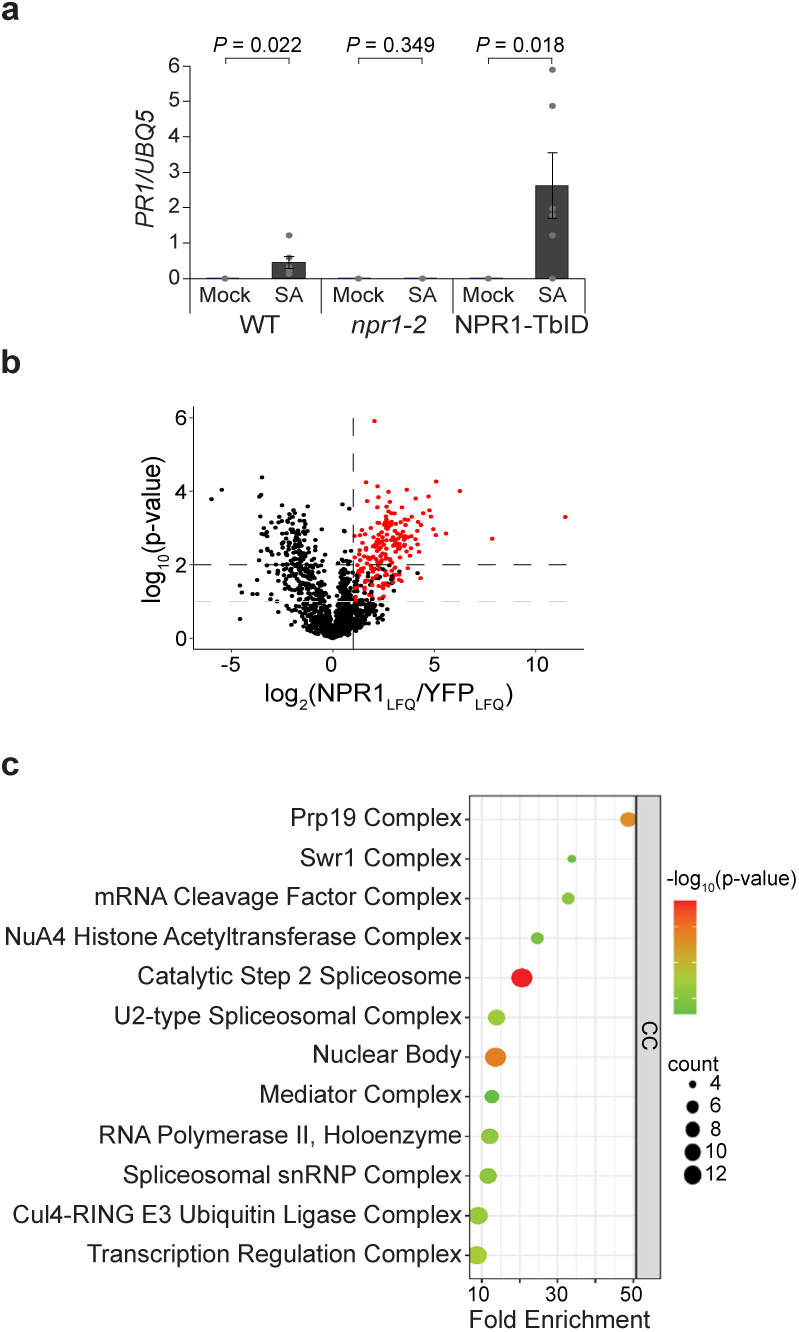
NPR1-TbID is biologically active and interacts with splicing and transcriptional machineries in the nucleus upon SA induction. **a**, *PR1* expression in WT, *npr1-2*, and *35S:NPR1-TbID/npr1-2* (*NPR1-TbID*) complementation plants treated with H_2_O (mock) or 1 mM SA for 24 h. (n = 6; error bars represent SEM, two-sided t-test was used to compare mock and 1 mM SA-treated samples). **b,** NPR1 proximal proteins 4 h after treatment with 1 mM SA detected through TurboID biotin affinity purification followed by Label Free Quantification (LFQ) Mass Spectrometry processed under harsh conditions (see Methods). Red points represent proteins that have a NPR1_LQF_/YFP_LFQ_ ≥ 2 and p-value < 0.1 in both stringent and harsh washing conditions (see Methods) or p-value < 0.01 in at least one washing condition. **c,** The enriched cellular components (CC) of the 234 NPR1 proximal proteins.

**Extended Data Fig. 2.**
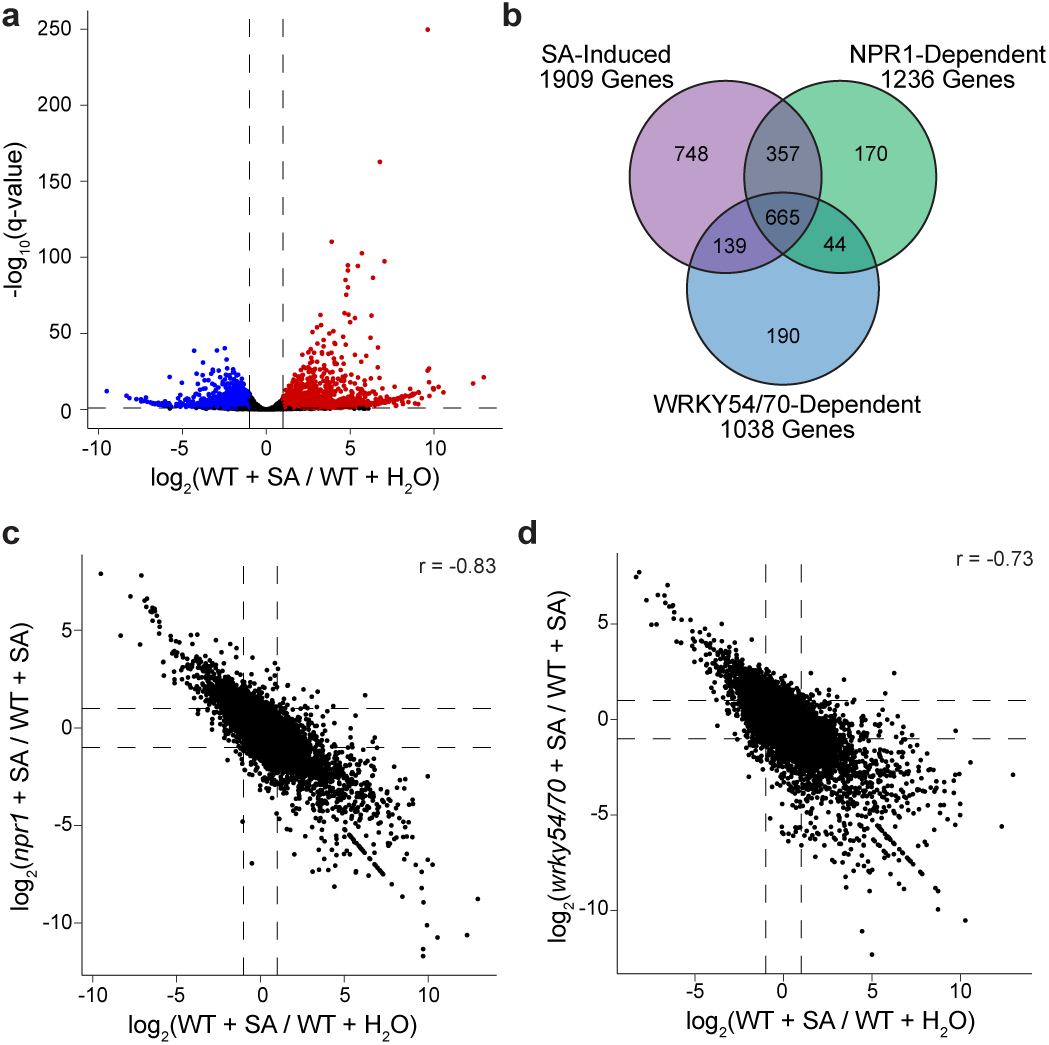
SA-mediated transcriptional changes are partially dependent on NPR1 and/or WRKY54/70. **a**, Volcano plot of SA-mediated transcriptional changes detected by QuantSeq. Colored points represent transcripts with (WT + SA) / (WT + H_2_O) ≥ 2 (red) or < -2 (blue) and an adjusted p-value (q-value) < 0.1. **b,** Venn diagram showing partial dependency of SA-mediated gene expression on NPR1 and/or WRKY54/70. **c, d,** Relationship between NPR1 (**c**) or WRKY70 (**d**) with SA-mediated transcriptional reprogramming. r, Pearson correlation coefficient.

**Extended Data Fig. 3.**
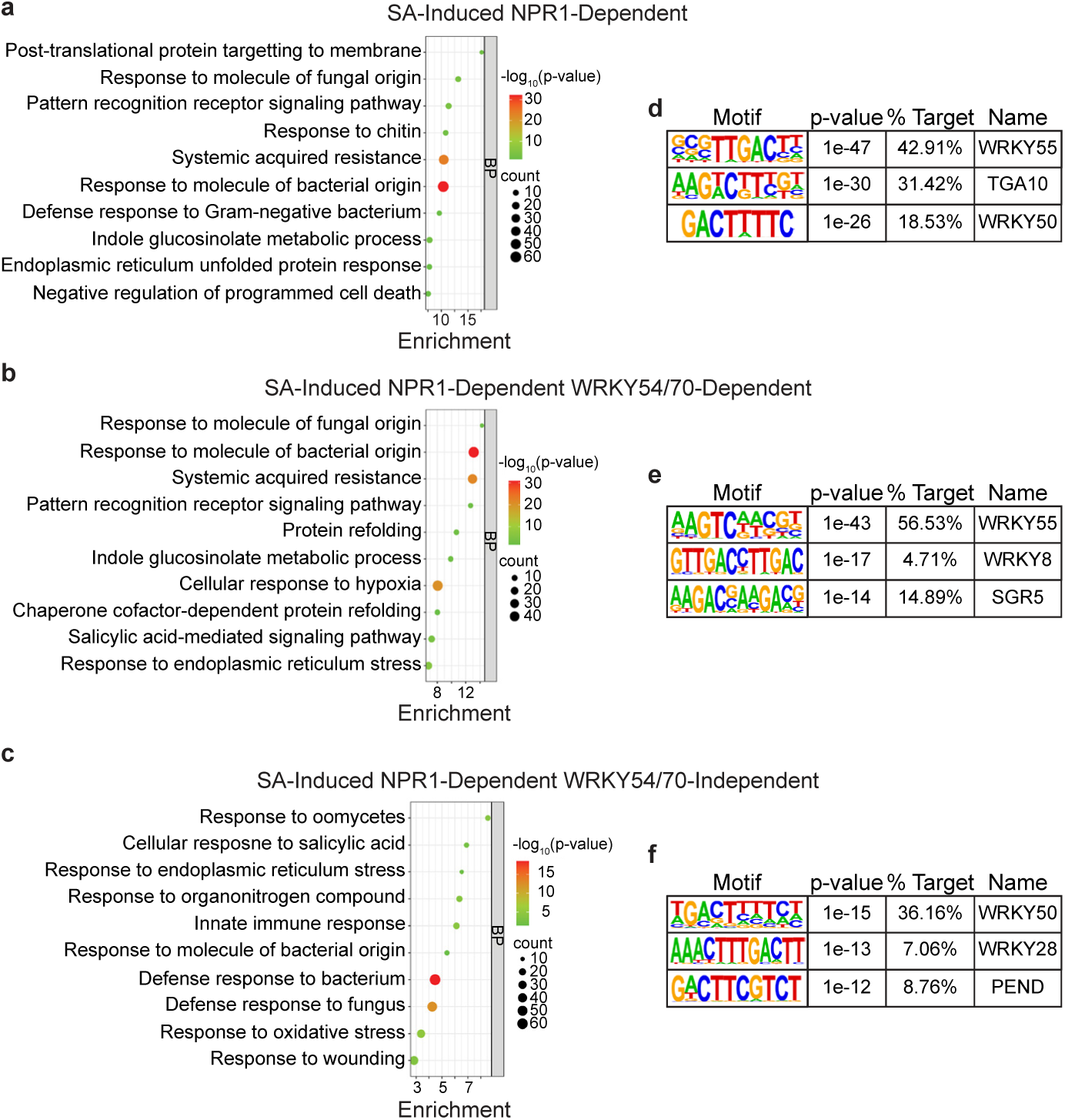
NPR1- and/or WRKY54/70-dependent genes are enriched in defense-related biological processes. **a-c**, Enriched biological processes (BP) in SA-induced NPR1-dependent genes (**a**), SA-induced NPR1-dependent WRKY54/70-independent genes (**b**), and SA-induced NPR1- and WRKY54/70-dependent genes (**c**). **d-f,** Motifs enriched from 1 kb upstream to 200 bp downstream of transcriptional start sites of the genes defined in **a-c**, respectively.

**Extended Data Fig. 4.**
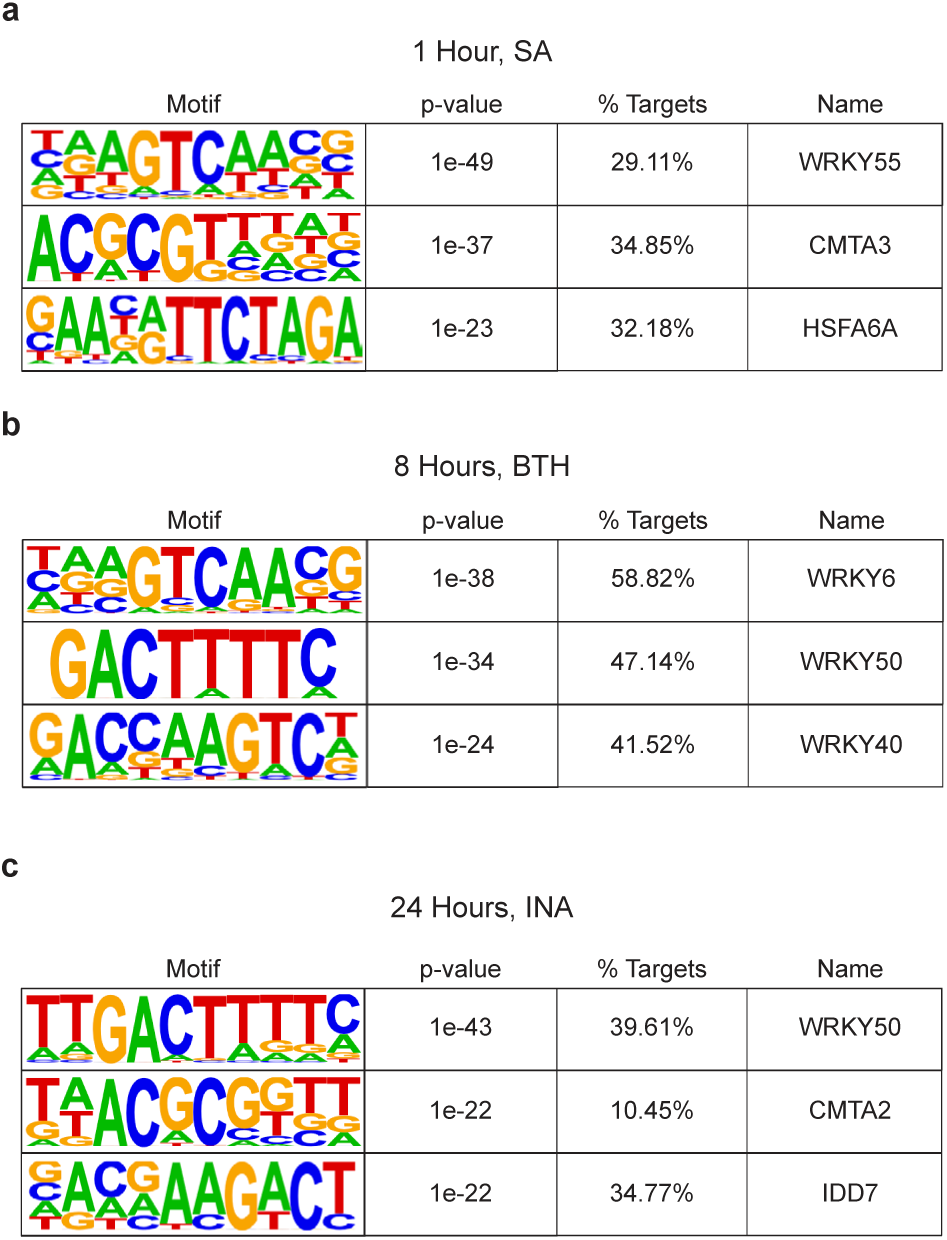
Motif enrichment of SA-, INA-, and BTH-induced genes from previously performed RNA-seq and microarray studies. **a**, Enriched motifs of SA-induced genes 1 h after treatment determined by RNA-seq^27^. **b,** Enriched motifs of the synthetic analog of SA, benzothiadiazole (BTH)-induced genes 8 h after treatment determined by microarray^2^. **c,** Enriched motifs of the synthetic analog of SA, 2,6-dichloroisonicotinic acid (INA)-induced genes 24 h after treatment determined by RNA-seq^16^.

**Extended Data Fig. 5.**
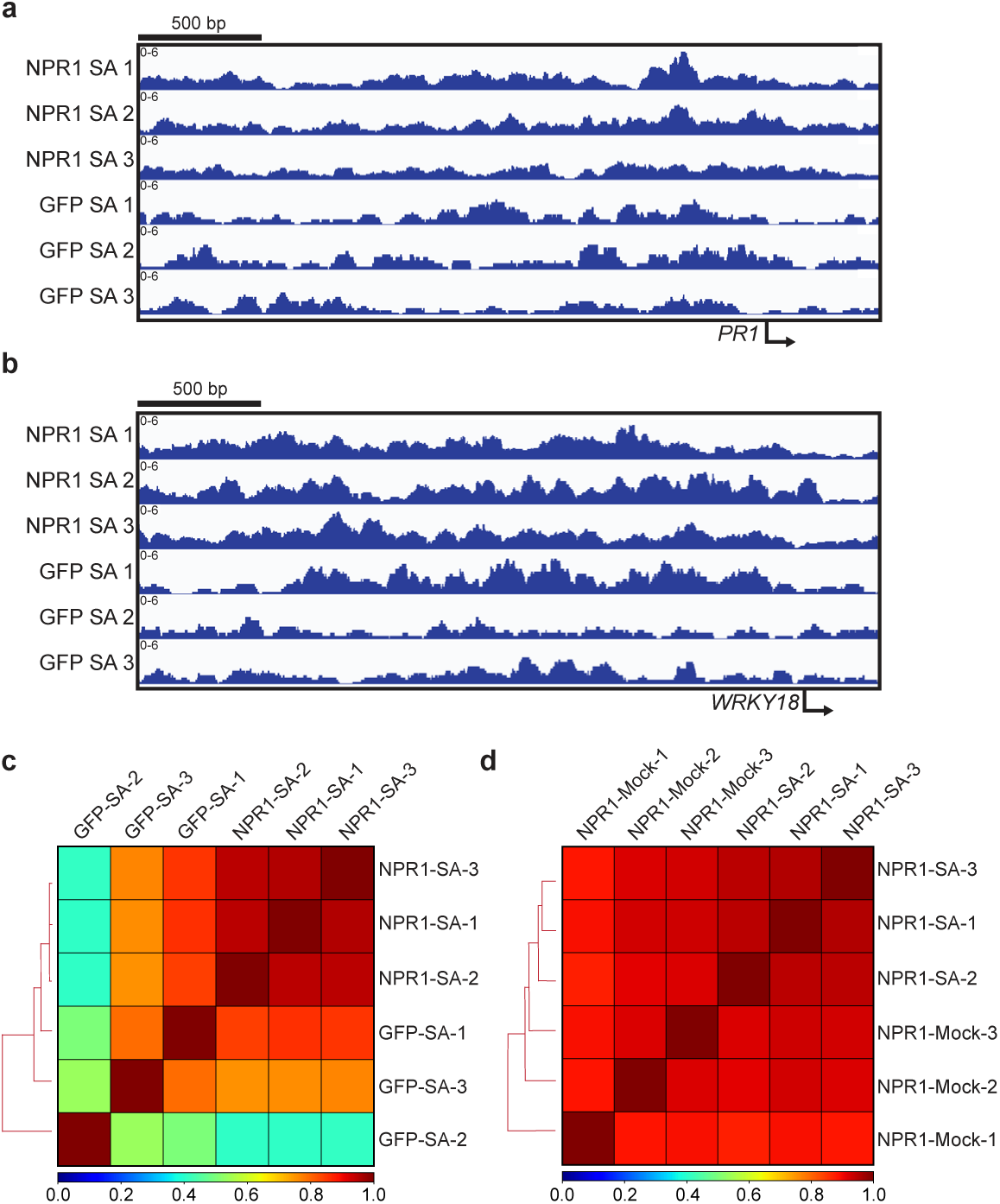
CUT&RUN failed to detect NPR1 binding to the chromatin in response to SA. **a, b**, Integrative Genomics Viewer (IGV) of normalized NPR1-GFP (NPR1) and GFP reads at the promoters of known NPR1-target genes *PR1* (**a**) and *WRKY18* (**b**) 4 h after 1 mM SA treatment. Numbers represent 3 biological replicates for each genotype. **c, d,** Pearson’s correlation between NPR1 and GFP treated with SA (**c**), between NPR1-GFP treated with H_2_O (mock) and SA (**d**).

**Extended Data Fig. 6.**
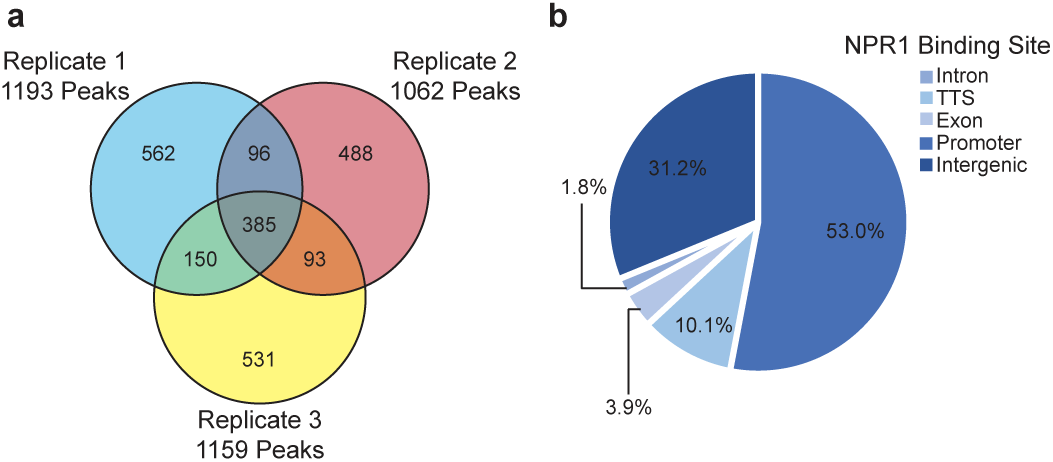
greenCUT&RUN detection of NPR1-GFP-binding at gene promoters upon SA induction. **a**, Venn diagram illustrating the reproducibility of greenCUT&RUN peaks among the three NPR1-GFP replicates 4 h after 1 mM SA treatment using GFP as the control. **b,** Pie chart illustrating the locations of NPR1-GFP peaks in its target genes defined as promoters (1 kb upstream to 1 bp upstream), intergenic (> 1 kb upstream), exon, intron, and transcriptional termination site (TTS).

**Extended Data Fig. 7.**
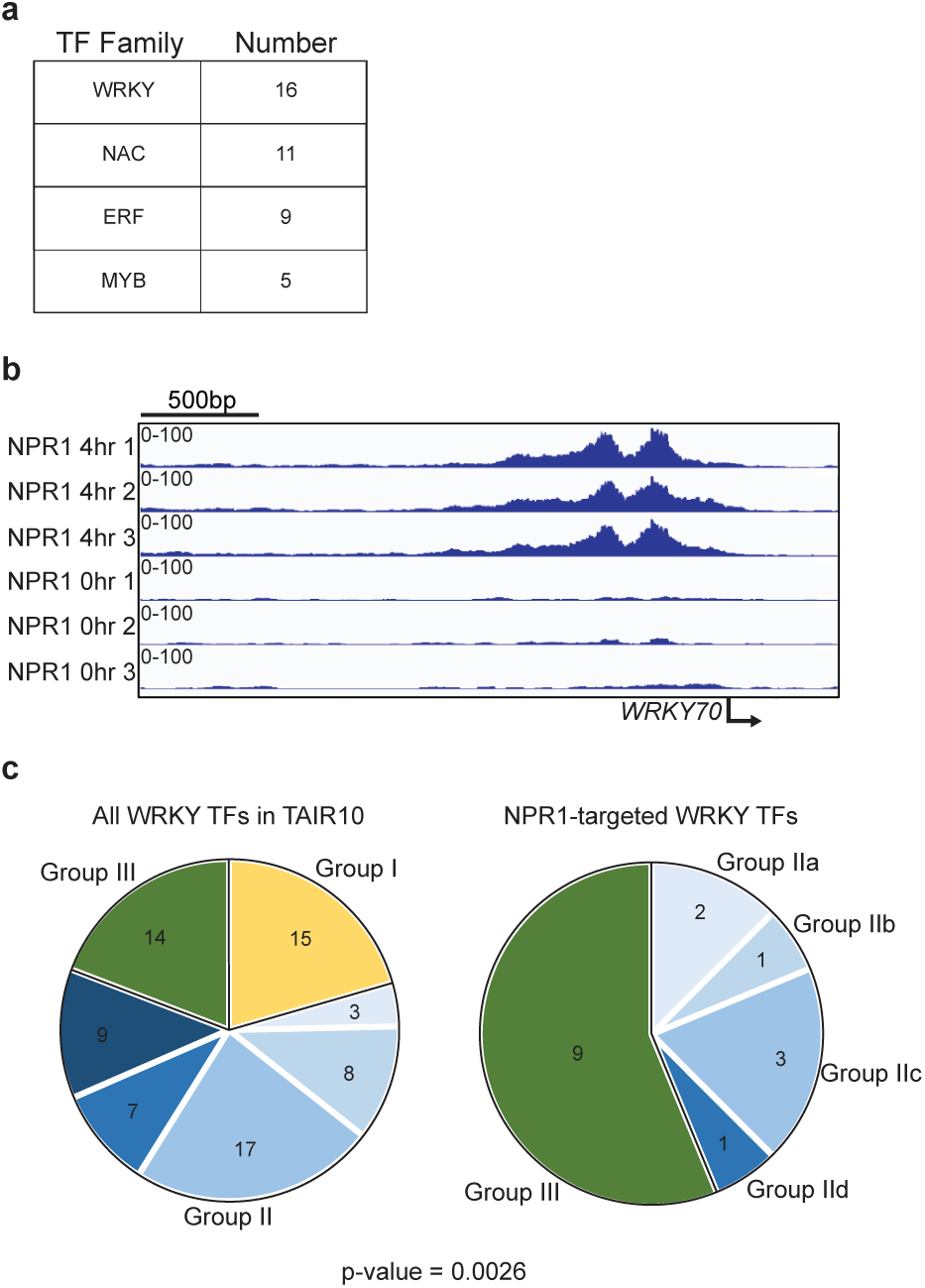
NPR1 predominantly targets Group III WRKY TFs. **a**, The most abundant TF families targeted by NPR1. **b,** Integrative Genomics Viewer (IGV) of normalized NPR1 reads with and without SA at the *WRKY70* promoter. Data from three biological replicates were used. **c,** Pie charts of all *Arabidopsis WRKY* TF genes based on The Arabidopsis Information Resource 10 (TAIR10) compared to *WRKY* genes directly targeted by NPR1 (statistical significance determined by chi-square test).

**Extended Data Fig. 8.**
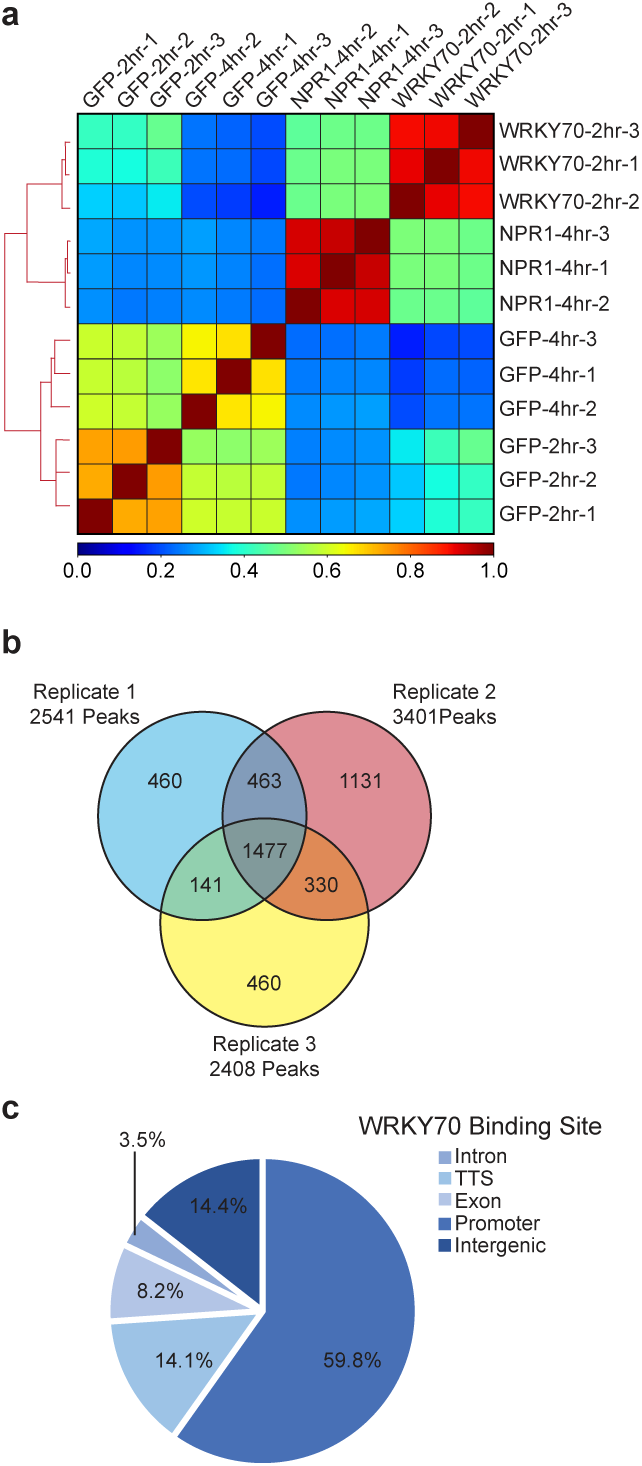
greenCUT&RUN detection of WRKY70-GFP-binding at gene promoters upon SA induction. **a**, Pearson correlation between WRKY70-GFP, NPR1-GFP, and GFP greenCUT&RUN data. **b,** Venn diagram illustrating the reproducibility of greenCUT&RUN peaks among the three WRKY70-GFP replicates. **c,** Pie chart illustrating the locations of WRKY70-GFP peaks in its target genes defined as promoters (1 kb upstream to 1 bp upstream), intergenic (> 1 kb upstream), exon, intron, and transcriptional termination site (TTS).

**Extended Data Fig. 9.**
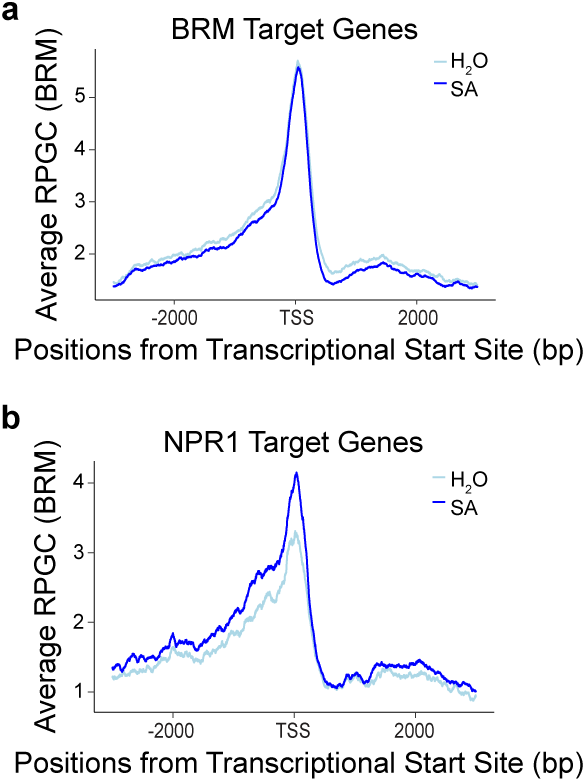
greenCUT&RUN of BRM-GFP with and without SA treatment. **a, b,** Mean profile of Reads Per Genomic Content (RPGC) of BRM-GFP reads 4 h after treatment with H_2_O or 1 mM SA at BRM-target genes (**a**) and NPR1-target genes (**b)**. TSS, transcriptional start site.

